# A nuclear export signal in KHNYN required for its antiviral activity evolved as ZAP emerged in tetrapods

**DOI:** 10.1101/2022.05.31.494266

**Authors:** Maria Jose Lista, Mattia Ficarelli, Harry Wilson, Dorota Kmiec, Rebecca L Youle, Joseph Wanford, Helena Winstone, Charlotte Odendall, Ian A Taylor, Stuart J D Neil, Chad M Swanson

## Abstract

The zinc finger antiviral protein (ZAP) inhibits viral replication by directly binding CpG dinucleotides in cytoplasmic viral RNA to inhibit protein synthesis and target the RNA for degradation. ZAP evolved in tetrapods and there are clear orthologs in reptiles, birds and mammals. When ZAP emerged, other proteins may have evolved to become cofactors for its antiviral activity. KHNYN is a putative endoribonuclease that is required for ZAP to restrict retroviruses. To determine its evolutionary path after ZAP emerged, we compared KHNYN orthologs in mammals and reptiles to those in fish, which do not encode ZAP. This identified residues in KHNYN that are highly conserved in species that encode ZAP, including several in the CUBAN domain. The CUBAN domain interacts with NEDD8 and Cullin-RING E3 ubiquitin ligases. Deletion of the CUBAN domain decreased KHNYN antiviral activity, increased protein expression and increased nuclear localization. However, mutation of residues required for the CUBAN domain-NEDD8 interaction increased KHNYN abundance but did not affect its antiviral activity or cytoplasmic localization, indicating that Cullin- mediated degradation may control its homeostasis and regulation of protein turnover is separatable from its antiviral activity. By contrast, the C-terminal residues in the CUBAN domain form a CRM1-dependent nuclear export signal (NES) that is required for its antiviral activity. Deletion or mutation of the NES increased KHNYN nuclear localization and decreased its interaction with ZAP. The final two positions of this NES are not present in fish KHNYN orthologs and we hypothesize their evolution allowed KHNYN to act as a ZAP cofactor.

**IMPORTANCE:** The interferon system is part of the innate immune response that inhibits viruses and other pathogens. This system emerged approximately 500 million years ago in early vertebrates. Since then, some genes have evolved to become antiviral interferon- stimulated genes (ISGs) while others evolved so their encoded protein could interact with proteins encoded by ISGs and contribute to their activity. However, this remains poorly characterized. ZAP is an ISG that arose during tetrapod evolution and inhibits viral replication. Because KHNYN interacts with ZAP and is required for its antiviral activity against retroviruses, we conducted an evolutionary analysis to determine how specific amino acids in KHNYN evolved after ZAP emerged. This identified a nuclear export signal that evolved in tetrapods and is required for KHNYN to traffic in the cell to interact with ZAP. Overall, specific residues in KHNYN evolved to allow it to act as a cofactor for ZAP antiviral activity.

## INTRODUCTION

Viral RNAs are targeted by diverse antiviral systems in eukaryotes to restrict viral replication. In plants and invertebrates, RNAi forms a major system to inhibit viral replication and this pathway was likely present in the last common ancestor of eukaryotes (1, 2). In vertebrates, the interferon system replaced RNAi as the predominant mechanism to inhibit viral replication, though RNAi is still active in some mammalian cell types, such as stem cells (1, 3). This protein-based system consists of pattern recognition receptors (PRRs) that bind pathogen-associated molecular patterns (PAMPs) (4). When the PRR binds a PAMP, signal transduction pathways are activated that induce transcription of antiviral cytokines, including interferons (IFNs) (5). The IFN system likely evolved in early vertebrates and is present in fish and the tetrapod lineages (amphibians, birds, reptiles, and mammals) (6). When IFNs are produced by an infected cell, they signal in an autocrine and paracrine manner to activate the expression of interferon-stimulated genes (ISGs). These have diverse functions and some are directly antiviral (7).

ZAP (also known as ZC3HAV1 or PARP13) is an ISG that evolved from a gene duplication of PARP12, which is an ISG expressed throughout the vertebrate lineage (8–11). ZAP emerged during tetrapod evolution and there are clear orthologs in mammals, birds and reptiles that have antiviral activity (8). ZAP binds viral RNA containing CpG dinucleotides to target it for degradation and inhibit its translation (11–15). ZAP interacts with several cellular proteins to form the ZAP antiviral system (16–22) and the emergence of ZAP may have provided an opportunity for these proteins to evolve in specific ways to act as cofactors, though little is known in this regard.

KHNYN is a ZAP-interacting protein that is required for it to restrict retroviruses (22). Unlike ZAP or its cofactor TRIM25, KHNYN is not a core mammalian ISG (9). It has two human paralogs, NYNRIN and N4BP1. NYNRIN evolved from a KHNYN gene duplication in which the RNase H and integrase domains from an endogenous retrovirus replaced the last exon of KHNYN (23). The function of this protein is unknown. N4BP1 is a predominantly nucleolar protein whose expression is induced by type I interferon (9, 24–27). While the specific functions of N4BP1 are still unclear, it has been shown to inhibit the NF-KB pathway as well as HIV-1 gene expression and the E3 ligase Itch (28–31). N4BP1 also has a genetic interaction with ZAP in that ZAP is required for N4BP1 antiviral activity (32).

Because it is unclear how KHNYN evolved to act as a ZAP cofactor, we characterized how specific changes in KHNYN that appeared after ZAP emerged in tetrapods regulate its antiviral activity. In mammalian and reptile species with clear ZAP orthologs, KHNYN contains a CRM1 nuclear export signal (NES) in its C- terminal CUBAN domain. This NES is required for KHNYN to traffic from the nucleus to the cytoplasm to interact with ZAP and mediate its antiviral activity. The penultimate and C-terminal positions of this NES evolved in the tetrapod lineage coincident with the emergence of ZAP.

## RESULTS

### Identification of residues in KHNYN that evolved coincident with the emergence of ZAP

While the complete N4BP1-KHNYN evolutionary pathway is currently unclear, N4BP1-like proteins are present in invertebrates such as echinoderms (phylum Echinodermata, e.g crown-of-thorns starfish) and molluscs (phylum Mollusca, e.g. California sea hare) (Fig. 1A and Fig. S1). These animals have one N4BP1-like protein. Within the phylum Chordata, N4BP1-like proteins are found in lancelets (subphylum Cephalochordata) and throughout the Vertebrata subphylum (Fig. 1A and Fig. S1). A single N4BP1-like protein is found in cartilaginous fishes (class Chondrichthyes), while bony fish (class Osteichthyes) and most tetrapods have clear N4BP1 and KHNYN orthologs including amphibians (class Amphibia), reptiles (class Reptilia) and mammals (class Mammalia). However, while N4BP1 orthologs are present in birds (class Aves), KHNYN orthologs are not, suggesting that it has been lost in this lineage.

**Fig. 1.**
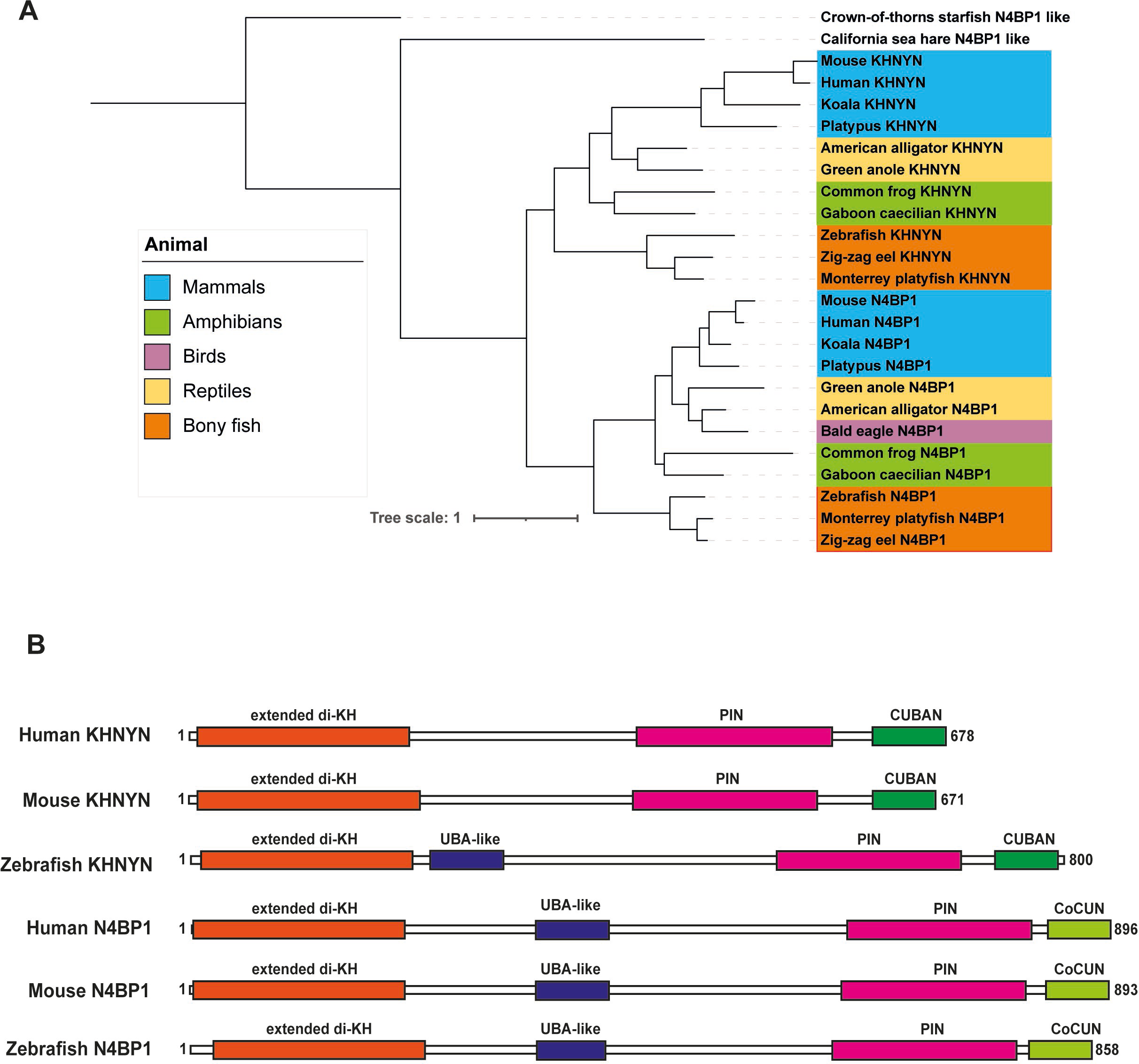
N4BP1 and KHNYN are paralogs in bony fish, amphibians, reptiles and mammals with a similar domain structure. **(A)** Maximum likelihood phylogenetic tree of KHNYN and N4BP1 amino acid sequences. Representative sequences from mammals (light blue), reptiles (yellow), birds (light purple), amphibians (green), and bony fish (orange) were aligned and maximum likelihood phylogeny was inferred using the LG substitution model DIVEIN. The crown-of-thorns starfish and Californian sea hare were used as outgroups to root the tree. An unpruned phylogenetic tree including all analyzed sequences, their scientific names and sequence accession numbers are presented in Figure S1. **(B)** Schematic of the extended di-KH, UBA-like, PIN and CUBAN/CoCUN domains predicted by AlphaFold in human, mouse, and zebrafish KHNYN and N4BP1.

N4BP1 has been reported to contain four structural domains: two N-terminal K homology (KH) domains in close proximity to form a di-KH domain, an ubiquitin-associated (UBA)-like domain, a PilT N-terminal (PIN) nuclease domain and a C-terminal Cousin of CUBAN (CoCUN) domain (24, 26, 29, 31, 33). Inspection of the AlphaFold models for human, mouse and zebrafish N4BP1 also supports the presence of these four structural domains (Fig. 1B, Fig. S2) (34, 35). The AlphaFold N4BP1 di-KH domain has three additional alpha helices at its C-terminus which is similar to a currently unpublished crystal structure for this domain (PDB 6q3v) and we refer to this as an extended di-KH domain (16). Because KHNYN had no known function before it was identified as a ZAP cofactor (22), it remains poorly characterized. We inspected the AlphaFold models for human, mouse and zebrafish KHNYN to determine its domain structure. The AlphaFold model for human and mouse KHNYN predicts three structural domains that match previously described domains: an extended di-KH domain, a PIN domain and a C-terminal cullin-binding domain associating with NEDD8 (CUBAN) domain (Fig. 1B and Fig. S2) (16, 26, 33- 36). It should be noted that the AlphaFold model for KHNYN needs to be experimentally validated and, of these three domains, only the CUBAN domain has been structurally characterized with three alpha helices present from residues 632- 678 (36).

To determine if the AlphaFold model for full-length KHNYN is supported by evolutionary conservation in mammalian KHNYN sequences, we used ConSurf (37, 38) to determine the normalized conservation score of each amino acid in a multiple sequence alignment (MSA) of placental mammals (Dataset S1) and plotted it against the structural confidence score (pLDDT) in the AlphaFold model of human KHNYN (Fig. 2A). Of note, the ConSurf conservation score corresponds to the evolutionary rate of the residue and lower numbers indicate more conserved residues. This analysis supports the three-domain model for mammalian KHNYN in that the regions with high AlphaFold confidence scores are well conserved evolutionarily, though an unstructured region from residues 339-359 also appears to be conserved (Fig. 2A and Dataset S1). The function of the extended di-KH domain in KHNYN is unknown but a deletion in this domain reduced its antiviral activity (22). The PIN domain in KHNYN is a putative endoribonuclease domain and mutation of potential catalytic residues in this domain inhibits its antiviral activity (22). The CUBAN domain binds NEDD8 and ubiquitin, both of which are members of the ubiquitin-like family, and preferentially binds monomeric NEDD8 over ubiquitin (36). NEDD8 binding mediates an interaction between KHNYN and neddylated cullin–RING E3 ubiquitin ligases (36). However, the role of the CUBAN domain for KHNYN antiviral activity is not known.

**Fig. 2.**
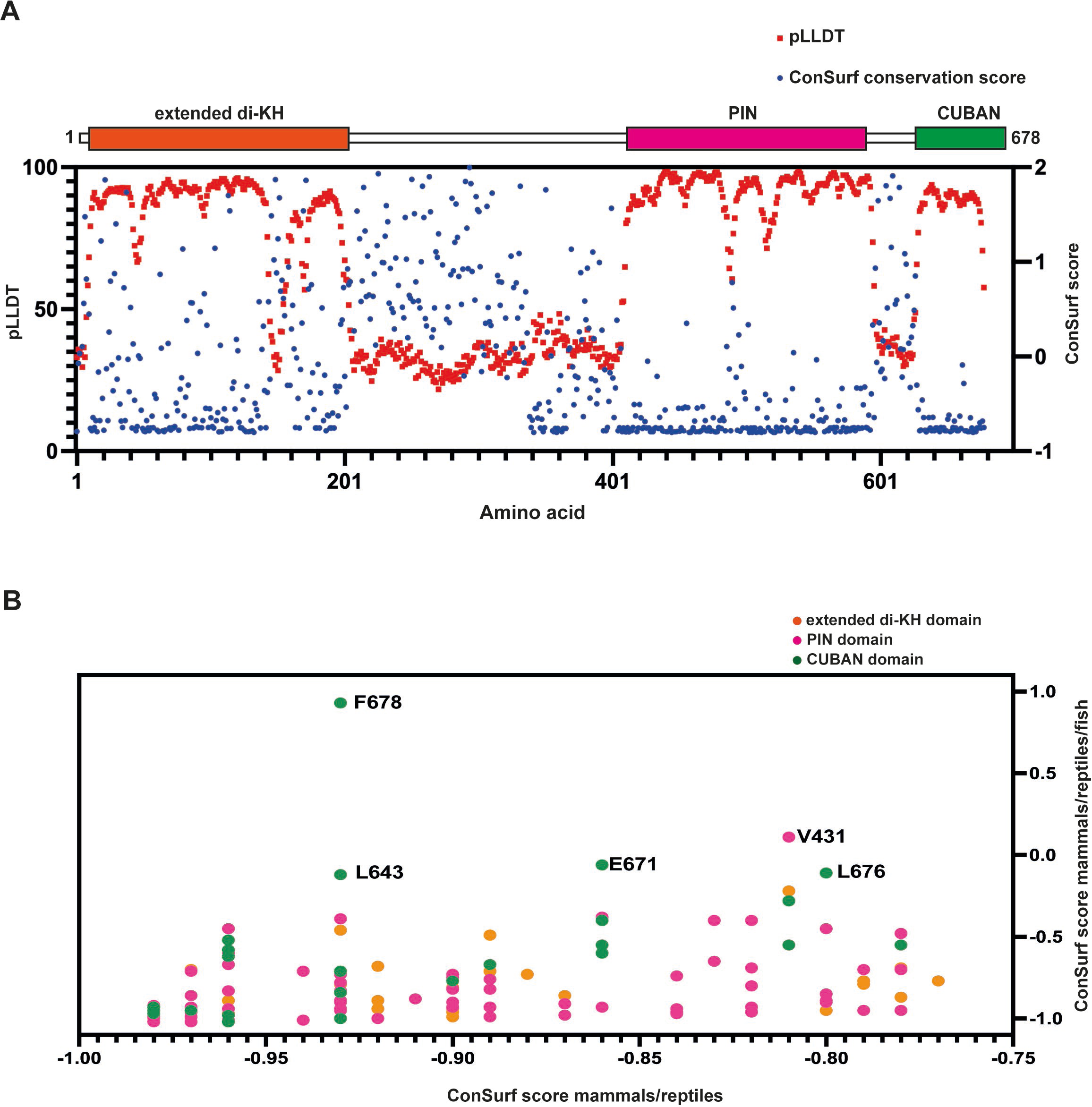
Some residues that are conserved in mammalian and reptile KHNYN orthologs are not conserved in bony fish orthologs. **(A)** KHNYN orthologs in placental mammals have three conserved domains. The AlphaFold structure confidence score (pLLDT) for human KHNYN and the ConSurf conservation score from a MSA of placental mammal KHNYN protein sequences were plotted for each amino acid in the human KHNYN sequence. A schematic showing the extended di-KH domain, PIN domain and CUBAN domain in KHNYN is shown above the plot. **(B)** Specific residues in KHNYN are less conserved in the mammal/reptile/bony fish MSA than the mammal/reptile MSA. The ConSurf conservation score for the mammal/reptile MSA (x-axis) was plotted against the ConSurf conservation score for the mammal/reptile/bony fish MSA (y-axis) for each residue in human KHNYN. Residues that substantially changed are labelled. Residues in the extended di-KH domain are colored orange, residues in the PIN domain are colored pink, and residues in the CUBAN domain are colored green.

The AlphaFold model for zebrafish KHNYN has five potential domains which are supported by evolutionary conservation in the bony fish lineage, including the extended di-KH domain, the PIN domain and the CUBAN domain (Fig. 1B, Fig. S2, Fig. S3, Dataset S1). One of the additional domains aligns with the UBA-like domain in N4BP1 and is not conserved in mammalian KHNYN orthologs (Fig. 1B, Fig. S2, Fig. S3) (31). The other potential domain has lower structure confidence scores than the other domains, does not appear to be present in mammalian KHNYN orthologs and the function of this region is unknown (Fig. S2 and Fig. S3). Overall, it appears that N4BP1 has four domains and these are conserved in bony fish KHNYN (Fig. 1B), indicating that this is the primordial domain structure for these proteins in vertebrates. The UBA-like domain has been lost in mammalian KHNYN orthologs and they have three clear structural domains.

To identify residues in KHNYN that evolved coincident with the emergence of ZAP, we used ConSurf to compare a MSA of mammal and reptile KHNYN orthologs (which encode a clear ZAP gene) and a MSA of mammal, reptile, and fish orthologs. ConSurf divides the conservation scores into nine categories. We used the residues in the category with the highest conservation (category 9, Dataset S1) in the mammal and reptile orthologs to compare the conservation scores for each residue in the mammal/reptile MSA to the score in the mammal/reptile/bony fish MSA (Fig. 2B). This identified five human residues (V431, L643, E671, L676 and F678) in which the conservation score substantially increased when fish were included in the MSA, indicating that these residues are not conserved in bony fish. Surprisingly, even though the CUBAN domain is the smallest domain in KHNYN, four of these five residues are in this domain. Since there is an experimentally determined structure for this domain, and its role for KHNYN antiviral activity is not known, we decided to focus on the CUBAN domain.

### The CUBAN domain in KHNYN regulates its abundance and subcellular localization

The CUBAN domain in KHNYN has been shown to bind to NEDD8 and mediate an interaction between KHNYN and neddylated cullin-RING E3 ubiquitin ligases, including CUL1, CUL2, CUL3 and CUL4 (36). However, it is not known whether the CUBAN domain is required for antiviral activity. Human immunodeficiency virus type 1 (HIV-1) is a common model system to study the antiviral activity of ZAP and its cofactors. While HIV-1 is highly depleted in CpG dinucleotides, which makes it poorly targeted by ZAP, when a specific region in HIV-1 *env* is engineered to contain additional CpGs through synonymous mutations (HIV-1_CpG_), the virus becomes ZAP-sensitive (15, 39–42). This allows matched ZAP-resistant and ZAP-sensitive viruses to be tested to characterize how ZAP and its cofactors restrict viral replication. To determine whether the CUBAN domain is required for KHNYN antiviral activity, we co-transfected either CRISPR-resistant wild-type pKHNYN or pKHNYN with a CUBAN domain deletion (KHNYNΔCUBAN, Fig. 3A) and either pHIV-1_WT_ or pHIV- 1_CpG_ into KHNYN CRISPR HeLa cells. Deletion of the CUBAN domain led to a large decrease in KHNYN antiviral activity on HIV-1_CpG_ infectious virus production, even though KHNYNΔCUBAN was expressed at much higher levels than the wild-type protein (Fig. 3B and Fig. S4A). This indicates that the CUBAN domain may be required for both KHNYN homeostatic turnover and antiviral activity.

**Fig. 3.**
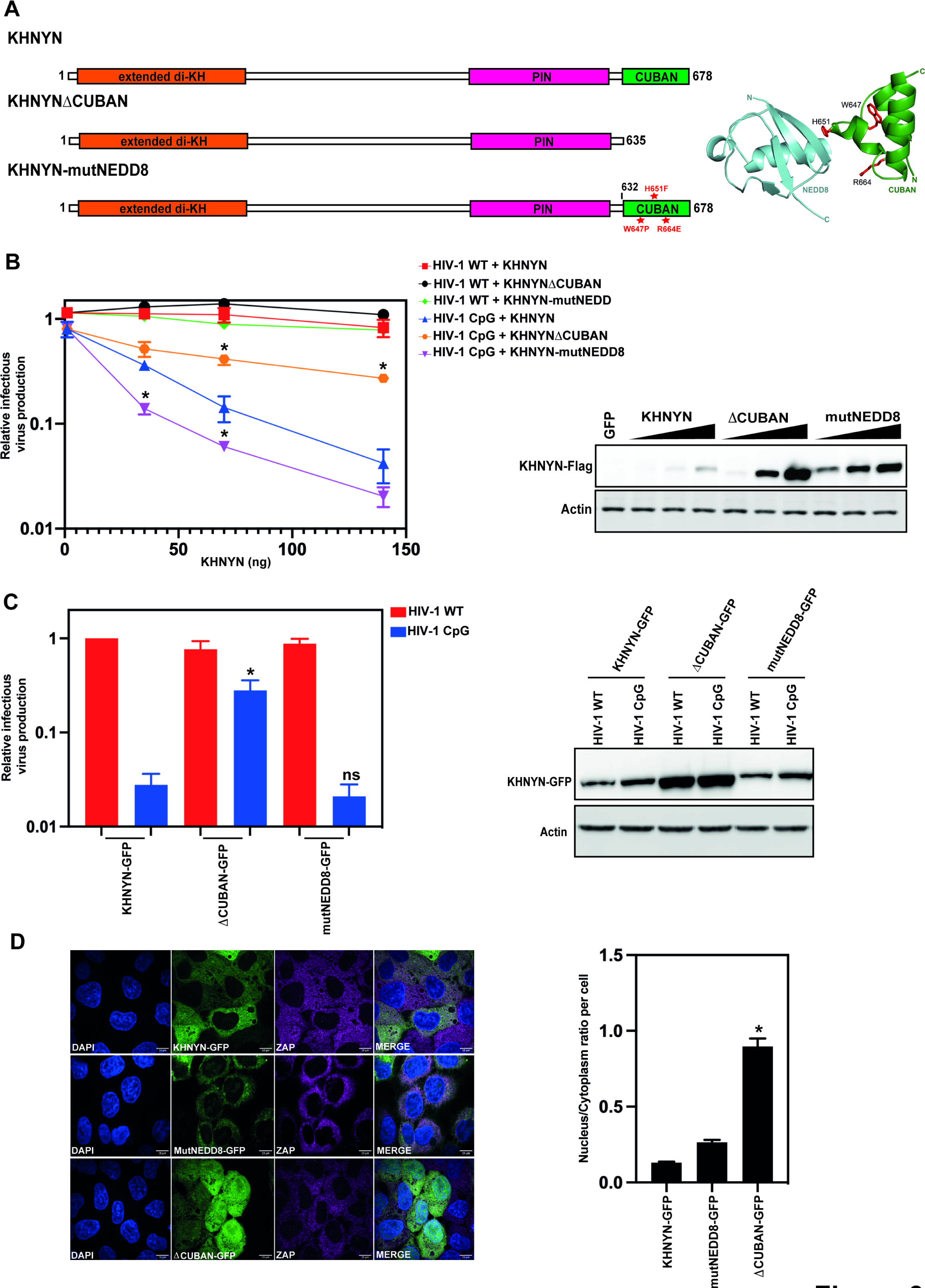
The CUBAN domain is essential for KHNHN antiviral activity and protein localization. **(A)** Left panel: Schematic representation of KHNYN, KHNYNΔ636-678 (KHNYNΔCUBAN) and KHNYN W647P/H651F/R664E (KHNYNmutNEDD8). The residues mutated in KHNYNmutNEDD8 are highlighted in red. Right panel: Cartoon representation of the NEDD8-CUBAN complex structure (PDB: 2N7K). The residues mutated in KHNYNmutNEDD8 are highlighted in stick representation in red. **(B)** Left panel: Infectious virus production from KHNYN CRISPR HeLa cells co-transfected with pHIV-1_WT_ or pHIV-1_CpG_ and increasing amount of CRISPR-resistant wild-type pKHNYN-Flag, pKHNYNΔCUBAN-Flag or pKHNYN-mutNEDD8-Flag plasmids. Each point shows the average value of three independent experiments normalized to the value obtained for pHIV-1_WT_ at 0 ng pKHNYN. *p < 0.05 as determined by a two- tailed unpaired t-test comparing wild-type KHNYN and the mutant KHNYN at each concentration in the HIV-1_CpG_ samples. Right panel: representative western blot of the protein level of wild type KHNYN-FLAG, KHNYNΔCUBAN-FLAG and KHNYNmutNEDD8-FLAG corresponding to the titration shown in the left panel. **(C)** Left panel: Infectious virus production from HeLa KHNYN CRISPR cells stably expressing wild-type KHNYN-GFP, KHNYNΔCUBAN-GFP or KHNYNmutNEDD8- GFP. All cell lines were transfected with HIV-1_WT_ or HIV-1_CpG_. Each bar shows the average value of five independent experiments normalized to the value obtained for wild type KHNYN co-transfected with pHIV-1_WT_. *p < 0.05 as determined by a two- tailed unpaired t-test comparing wild-type KHNYN and the mutant KHNYN construct in the HIV-1_CpG_ samples. Data are represented as mean ± SD. Right panel: Representative western blot for GFP showing the KHNYN-GFP protein levels in the wild-type KHNYN-GFP, KHNYNΔCUBAN-GFP or KHNYNmutNEDD8-GFP cell lines. **(D)** Left panel: Confocal microscopy images of the KHNYN-GFP cell lines (green) co-stained with endogenous ZAP (magenta), scale bar is 10µm. Right panel: Signal quantification per cell (50 cells total per condition) of the ratio of KHNYN nuclear and cytoplasmic distribution in the KHNYN-GFP cell lines. *p < 0.05 as determined by a two-tailed unpaired t-test comparing the nuclear/cytoplasmic ratio between each sample.

The CUBAN domain comprises a three-helix bundle (α1 = T632-R640, α2 = K652- L657, α3 = Y662-F678) connected by two loops (36). The binding interface between the CUBAN domain and NEDD8 is formed from a negatively charged motif in NEDD8 and a positively charged surface in the CUBAN α2 and surrounding residues. Quantitative proteomics in HeLa cells have shown that many of the ZAP cofactors required for viral RNA degradation are expressed at much higher levels than KHNYN (Fig. S4B) and there are ∼25-fold more ZAP molecules/cell than KHNYN (43). Therefore, the interaction between the CUBAN domain and NEDD8 could contribute to the low expression level of endogenous KHNYN (43). Three mutations have previously been shown to decrease binding of the CUBAN domain to NEDD8: W647P, H651F and R664E (Fig. 2A) (36). When these mutations were introduced in KHNYN (KHNYN-mutNEDD8), they increased its protein abundance and moderately increased its antiviral activity for HIV-1_CpG_ (Fig. 3B and Fig. S4A). This suggests that the CUBAN domain-NEDD8 interaction likely regulates KHNYN turnover but not antiviral activity and indicates that there may be another functional motif in the CUBAN domain.

To analyze the subcellular localization of the mutant KHNYN proteins, KHNYN-GFP, KHNYN-mutNEDD8-GFP and KHNYNΔCUBAN-GFP were stably expressed in the KHNYN CRISPR HeLa cells. As expected, the KHNYN-GFP cells restricted HIV-1_CpG_ (Fig. S5A). To confirm the effect of the CUBAN domain mutations in the context of the KHNYN-GFP stable cell lines, they were transfected with pHIV-1_WT_ or pHIV-1_CpG_.

Similar to the transient transfection experiments described above (Fig. 3B), deletion of the CUBAN domain decreased KHNYN-GFP antiviral activity while introducing the mutations that reduce NEDD8 binding did not affect it (Fig. 3C and Fig. S5B). Of note, the increase in KHNYN abundance due to the CUBAN domain deletion or mutations that decrease NEDD8 binding are less pronounced in the KHNYN-GFP cell lines than in the experiments with transiently transfected KHNYN-FLAG constructs (Fig. 3C), possibly because the GFP fusion stabilizes the wild-type protein. Interestingly, while KHNYN-mutNEDD8-GFP had a similar localization to the cytoplasm as wild-type KHNYN-GFP, KHNYNΔCUBAN-GFP had a substantial increase in nuclear localization (Fig. 3D). Therefore, the CUBAN domain regulates KHNYN subcellular localization in addition to its homeostatic turnover and antiviral activity.

### KHNYN has a nuclear export signal at the C-terminus of the CUBAN domain that is required for antiviral activity

CRM1 (also known as XPO1) is a nuclear export protein that mediates trafficking of many cellular proteins and ribonucleoprotein complexes from the nucleus to the cytoplasm (44). CRM1 binds leucine-rich nuclear export signals (NESs) in cargo proteins and KHNYN has previously been identified as a CRM1 cargo in a large-scale proteomics screen (44, 45). To confirm that KHNYN uses the CRM1 nuclear export pathway, we compared the subcellular localization of KHNYN-GFP, KHNYN- mutNEDD8-GFP and KHNYNΔCUBAN-GFP in the absence and presence of leptomycin B, a small molecule inhibitor of CRM1 (46). Addition of leptomycin B to the KHNYN-GFP stable cell lines substantially increased wild-type KHNYN-GFP or KHNYN-mutNEDD8-GFP nuclear localization to levels similar to KHNYNΔCUBAN-GFP (Fig. 4). However, the subcellular localization of KHNYNΔCUBAN-GFP was not affected by leptomycin B, indicating that the CRM1 NES is present in the CUBAN domain. While ZAP also has a CRM1 NES and has been reported to be a CRM1- dependent nucleocytoplasmic shuttling protein (45, 47), it was not substantially re- localized to the nucleus by leptomycin B treatment (Fig. 4). This suggests that, in these cells, ZAP is sequestered in the cytoplasm and is not undergoing CRM1- dependent nuclear-cytoplasmic trafficking.

**Fig. 4.**
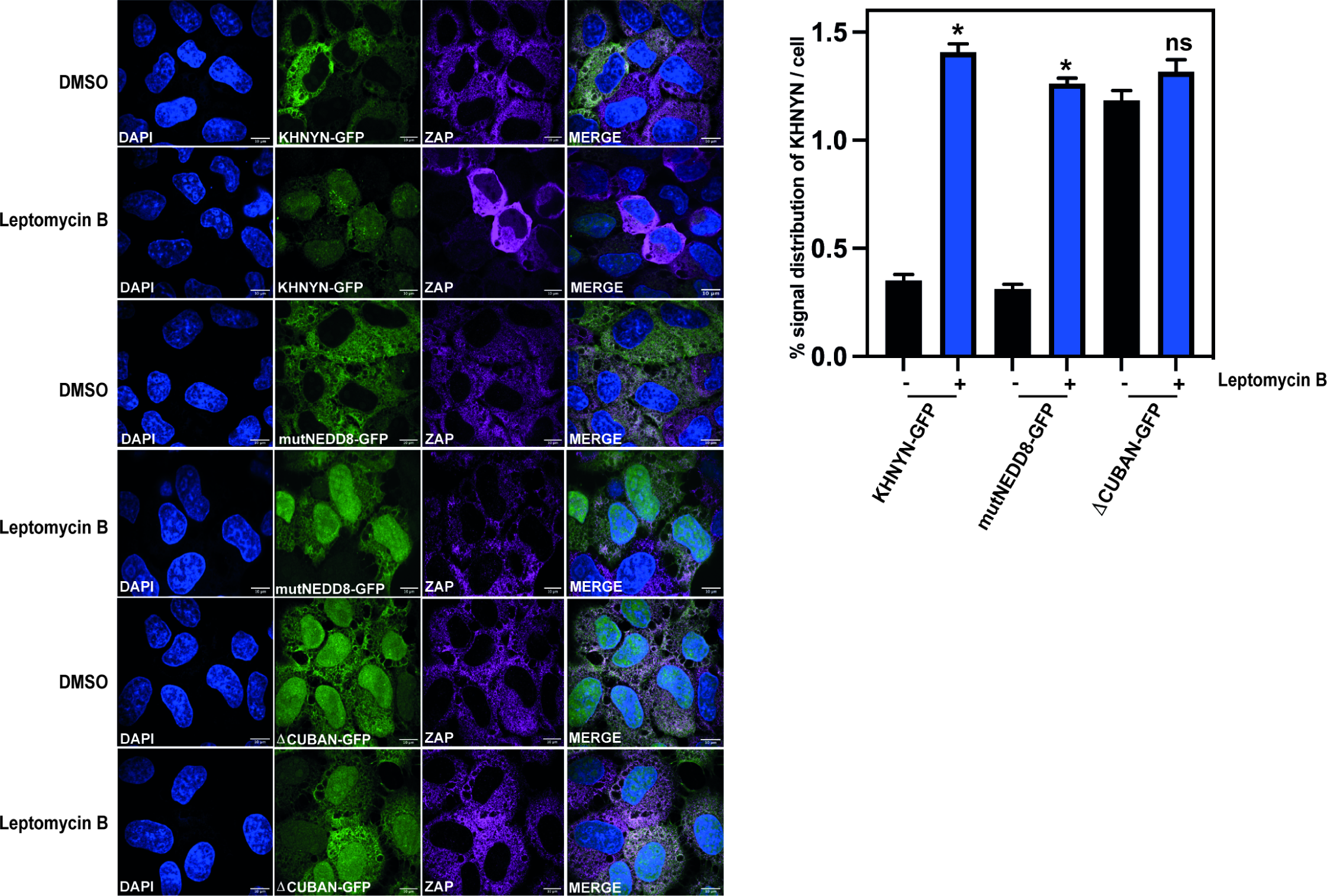
CRM1 inhibition by Leptomycin B re-localizes KHNYN to the nucleus. Left panel: Representative confocal microscopy images of KHNYN CRISPR HeLa cells stably expressing either wild-type KHNYN-GFP or the indicated mutants before and after a four-hour treatment with 50 nM Leptomycin B. KHNYN-GFP is shown in green, endogenous ZAP co-staining is shown in magenta, scale bar is 10 µm. Right panel: Signal quantification per cell (50 cells in total per condition) of the ratio of KHNYN nuclear and cytoplasmic distribution for KHNYN-GFP or the indicated mutant protein before and after Leptomycin B treatment. *p < 0.05 as determined by a two-tailed unpaired t-test comparing the nuclear/cytoplasmic ratio between each sample.

There are at least two types of NESs in CRM1 cargo proteins with different spacing of hydrophobic residues that fit into five pockets in CRM1: the Rev-type NES and the PKI-type NES (48). To identify potential NESs in KHNYN, we used the Wregex tool (49), which identified a putative NES at the C-terminus of the CUBAN domain. This NES has hydrophobic residues with the PKI-type NES spacing (residues 669-678, <u>L</u>SEA<u>L</u>LS<u>L</u>N<u>F</u>, amino acids predicted to bind CRM1 are underlined). Of note, this tool only identified positions 1-4 for the predicted CRM1 NES and did not identify the more recently identified position 0 (48, 49). For a PKI-type NES, position 0 is two amino acids upstream of position 1 and is preferentially preceded by an acidic residue (48). The C-terminal NES in KHNYN fits this consensus perfectly, with the full NES predicted to be DINQ<u>L</u>SEA<u>L</u>LS<u>L</u>N<u>F</u> (Fig. 5A). This sequence is located in the third helix of the CUBAN domain (Fig. 5B), which does not contain any of the residues that interact with NEDD8.

**Fig. 5.**
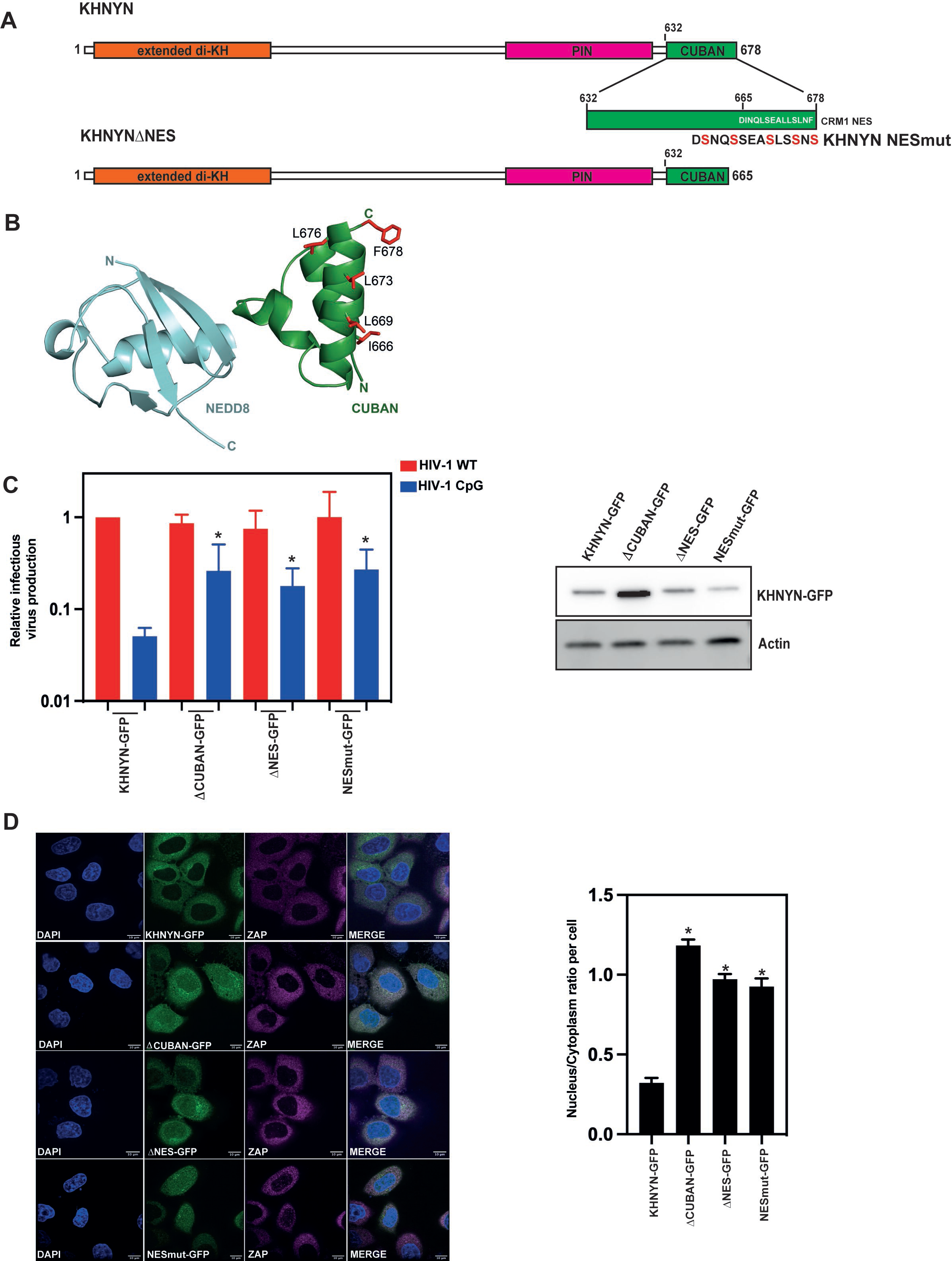
A nuclear export signal present in the CUBAN domain is required for KHNYN cytoplasmic location and antiviral activity. **(A)** Schematic representation of the KHNYN CUBAN domain and nuclear export signal. The residues that were mutated in KHNYN-NESmut are shown in red. **(B)** Cartoon representation of the NEDD8-CUBAN complex structure (PDB: 2N7K). Residues in the NES that were mutated to serine are shown as sticks in red. **(C)** Left panel: Infectious virus production from KHNYN CRISPR HeLa cells stably expressing wild-type KHNYN- GFP, KHNYNΔCUBAN-GFP, KHNYNΔNES-GFP or KHNYN-NESmut-GFP transfected with HIV-1_WT_ or HIV-1_CpG_. Each bar shows the average value of three independent experiments normalized to the value obtained for wild type KHNYN co- transfected with pHIV-1_WT_. *p < 0.05 as determined by a two-tailed unpaired t-test comparing each mutant KHNYN-GFP sample to wild type KHNYN-GFP. Right panel: Representative western blot showing KHNYN-GFP protein levels. **(D)** Left panel: Confocal microscopy images of the KHNYN-GFP cell lines (green) co-stained for endogenous ZAP (magenta), scale bar is 10 µm. Right panel: Signal quantification per cell (50 cells total per condition) of the ratio of KHNYN nuclear to cytoplasmic distribution in the KHNYN-GFP cell lines. *p < 0.05 as determined by a two-tailed unpaired t-test comparing the nuclear/cytoplasmic ratio between each sample.

To test the functional role of the NES in KHNYN, we made stable cell lines expressing KHNYNΔNES-GFP and KHNYN-NESmut-GFP. KHNYN-NESmut-GFP has all five amino acids predicted to directly bind CRM1 mutated to serine and, in KHNYNΔNES-GFP, the NES sequence was deleted (Figure 5A). Deleting or mutating the NES decreased KHNYN antiviral activity and increased its nuclear localization similar to KHNYN-ΔCUBAN (Fig. 5C-D and Fig. S6A). This suggests that the loss of antiviral activity for KHNYNΔCUBAN is due to the deletion of the C-terminal NES in the CUBAN domain. While deletion or mutation of the NES localizes KHNYN to the nucleus, it has no effect on ZAP or TRIM25 localization (Fig. 5D and Fig. S6B).

### Evolution of the KHNYN CUBAN domain

To further analyze how the CUBAN domain in KHNYN has evolved since ZAP emerged, we calculated the ratio of the percentage of the dominant amino acid at a specific site in the mammal/reptile MSA to the percentage of the dominant residue at that site in the mammal/reptile/bony fish MSA (see materials and methods for more detail, Dataset S1). We then plotted the ratio for the highly conserved residues in the in the mammal/reptile MSA (Fig. 6A, categories 7, 8 and 9 in the ConSurf analysis, Dataset S1). Two clear distributions of the amino acids are present: residues that are highly conserved in KHNYN from bony fish to mammals (ratio >0.85, colored in cyan) and residues that are conserved only in the mammal and reptile lineage (ratio <0.75, colored in magenta) (Fig. 6A). Inspection of the CUBAN domain structure (36) shows that many of the highly conserved residues in mammals, reptiles and bony fishes are likely essential for the overall fold including T365, L638, R639, F646, G648, V653, L657, P661, N667 and L669. Many of these residues are also conserved in cartilaginous fishes, lampreys, lancelets as well as invertebrates (echinoderms and molluscs) (Fig. S7), highlighting that a CUBAN-like domain is an ancestral feature of the N4BP1-like proteins.

**Fig. 6.**
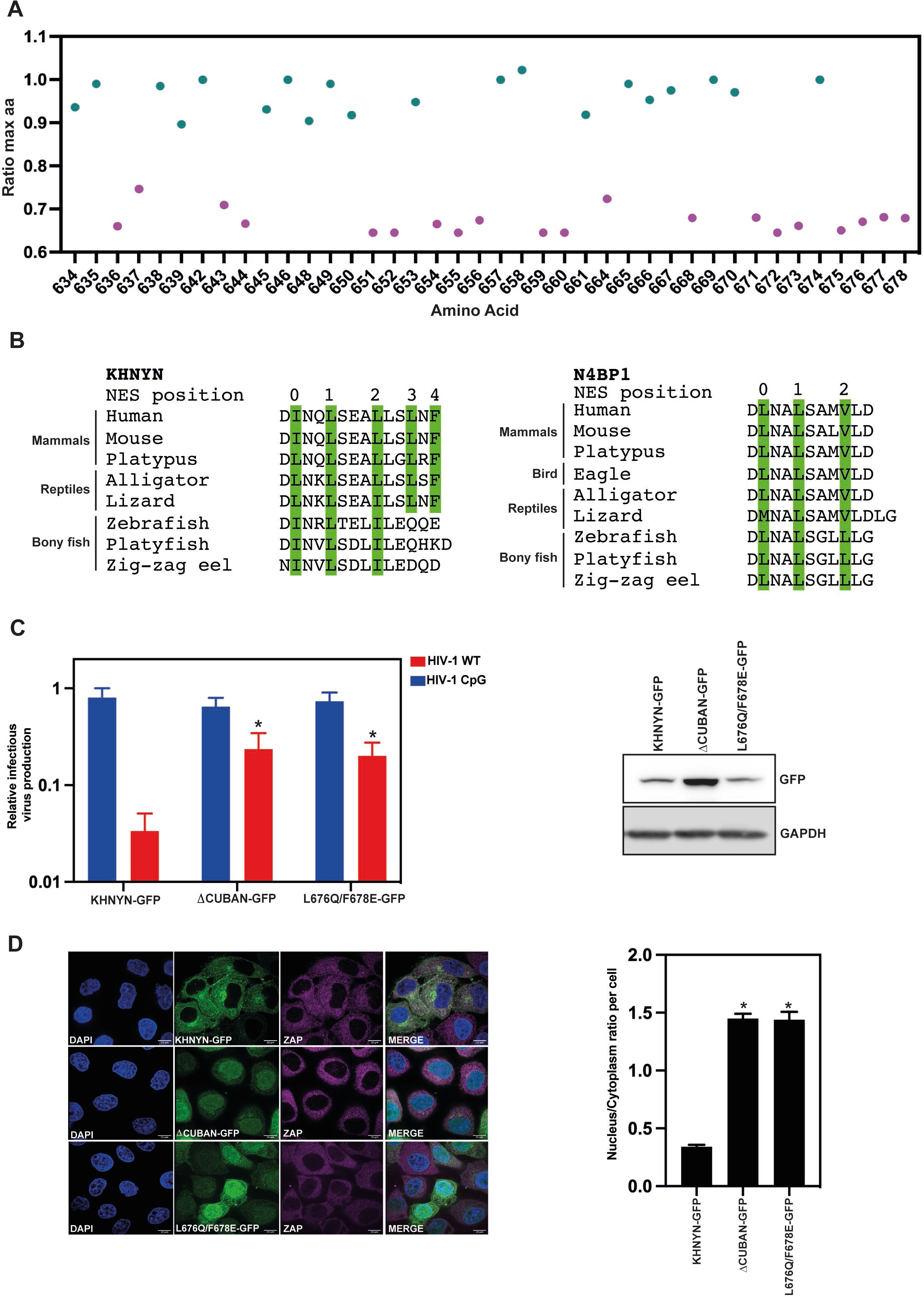
Positions 3 and 4 in the KHNYN NES in the CUBAN domain evolved at a similar time as ZAP in tetrapods and are required for KHNYN antiviral activity and cytoplasmic localization. (A) Ratio of the percentage of the maximum amino acid present at each position in the CUBAN domain in the mammal/reptile MSA to the mammal/reptile/bony fish MSA. Residues with a ratio >0.85 are colored cyan and residues with a ratio <0.75 are colored magenta. Residues that were not highly conserved in the mammal/reptile MSA (ConSurf categories 7, 8 and 9) are not shown. (B) Alignment of the C-terminal NES residues of KHNYN and N4BP1 from selected species. Residues in PKI-like NES positions 0 – 4 are highlighted in green. (C) Left panel: Infectious virus production from KHNYN CRISPR HeLa cells stably expressing wild-type KHNYN-GFP, KHNYNΔCUBAN-GFP or KHNYN L676Q/F678E transfected with HIV-1_WT_ or HIV-1_CpG_. Each bar shows the average value of three independent experiments normalized to the value obtained for wild type KHNYN co- transfected with pHIV-1_WT_. *p < 0.05 as determined by a two-tailed unpaired t-test comparing each mutant KHNYN-GFP sample to wild type KHNYN-GFP. Right panel: Representative western blot showing KHNYN-GFP protein levels. (D) Left panel: Confocal microscopy images of the KHNYN-GFP cell lines (green) co-stained for endogenous ZAP (magenta), scale bar is 10µm. Right panel: Signal quantification per cell (50 cells total per condition) of the ratio of KHNYN nuclear to cytoplasmic distribution in the KHNYN-GFP cell lines. *p < 0.05 as determined by a two-tailed unpaired t-test comparing nuclear/cytoplasmic ratio between each sample.

Several of the residues that are highly conserved in mammal and reptile KHNYN orthologs but not those in bony fish are part of the basic binding site in the CUBAN domain that interacts with NEDD8, including H651 and K652. In addition, many of the residues that make up the NES are conserved in mammals and reptiles but not in bony fish. Looking at the five residues in the NES that interact with CRM1, interesting patterns emerge. I666 and L669 at positions 0 and 1 in the NES are highly conserved in KHNYN orthologs and are also conserved as a bulky hydrophobic residue in N4BP1 and N4BP1-like proteins (Fig. 6B and Fig. S7). Isoleucine and leucine are the strongest amino acids for a PKI-type NES at positions 0 and 1 (48). At position 2 in the NES, a bulky hydrophobic residue is also often present in KHNYN and N4BP1 orthologs (Fig. 6B and Fig. S7). However, it is typically a leucine in tetrapod KHNYN orthologs in contrast to bony fish KHNYN orthologs, N4BP1 or N4BP1-like proteins, where it is usually an isoleucine or valine, both of which are weaker in the context of a PKI-type NES (48). L676 and F678 at positions 3 and 4 in the NES were among the top five residues with the largest differential in the ConSurf analysis comparing the mammal/reptile KHNYN MSA to the mammal/reptile/bony fish MSA (Fig. 2B). L676 is not present in most bony fish KHNYN orthologs, N4BP1 orthologs and N4BP1-like proteins. (Fig. 6B and Fig. S7). F678 is not found in any of the sequences analyzed outside of tetrapod KHNYN orthologs. To determine whether the NES is functional without leucine and phenylalanine at positions 3 and 4, we mutated L676 and F678 to the residues found in zebrafish KHNYN (L676Q, F676E) and made stable cell lines expressing these proteins with a GFP tag. As expected, KHNYN L676Q/F678E-GFP had a substantial loss of antiviral activity and was localized in the nucleus (Fig. 6C-D, Fig. S8). This indicates that the tetrapod-specific changes in the C-terminus of KHNYN created a NES that allows it to act as a ZAP cofactor.

### The nuclear export signal in the CUBAN domain is required for KHNYN to interact with ZAP-L

There are two predominant isoforms for ZAP, ZAP-L and ZAP-S (11, 50). ZAP-L contains a C-terminal S-farnesylation modification that localizes it to the cytoplasmic endomembrane system while ZAP-S appears to be diffuse in the cytosol (51–54). ZAP-L has greater antiviral activity than ZAP-S against some viruses, including HIV- 1_CpG_, and this requires the S-farnesylation post-translational modification (50–53). ZAP-L interacts with KHNYN more efficiently than ZAP-S and the interaction between ZAP-L and KHNYN is decreased when the CaaX box in ZAP-L that mediates S-farnesylation is mutated (53). Therefore, the NES in KHNYN may be necessary for its antiviral activity because it is required for KHNYN to traffic to the cytoplasm so it can interact with ZAP-L. To determine if the nuclear export signal in KHNYN is required for it to interact with ZAP, we performed co-immunoprecipitation experiments in the KHNYN-GFP, KHNYNΔNES-GFP and KHNYN L676Q/F678E- GFP cell lines. Immunoprecipitation of KHNYN-GFP pulled down ZAP-L but little ZAP-S (Fig. 7). KHNYNΔNES-GFP and KHNYN L676Q/F678E-GFP immunoprecipitated less ZAP-L than wild-type KHNYN (Fig. 7), which suggests that nuclear export of KHNYN is required for it to interact with ZAP-L localized to the cytoplasmic endomembrane system.

**Fig. 7.**
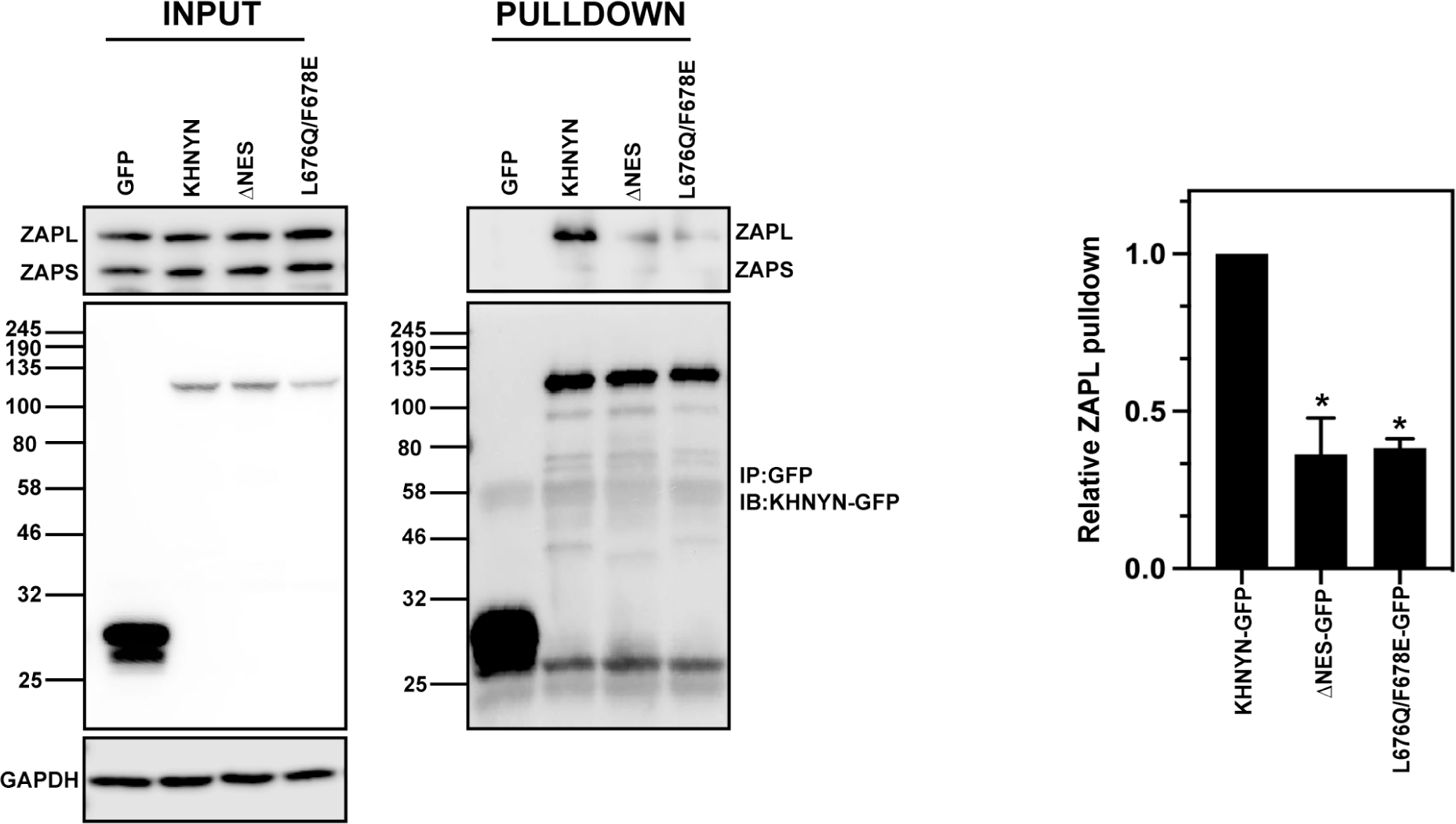
The NES in the KHNYN CUBAN domain promotes its interaction with ZAP. Left panel: GFP control, KHNYN-GFP and KHNYNΔNES-GFP HeLa cells were lysed and immunoblotted for GFP and endogenous ZAP. Middle panel: GFP was immunoprecipitated in each cell lysate and blotted for GFP or endogenous ZAP. Right panel: The amount of ZAP-L immunoprecipitated relative to the KHNYN-GFP sample is presented in a bar graph, N = 3. * p = <0.05 as determined by a two-tailed unpaired t-test.

## DISCUSSION

The development of the interferon system has led to extensive evolution in many genes (55). Many ISGs have been shown to have undergone periods of rapid evolution and a panel of ISGs have been shown to evolve faster than interferon-induction genes or a panel of random genes (56). However, little is known about the evolution of non-ISGs that encode ISG cofactors and how these genes evolve after the emergence of an ISG. Importantly, this evolutionary signature is likely to be different from the signature induced by a viral countermeasure to an ISG. Instead of a positive natural selection signature, evolutionary changes that allow a protein to act as an ISG cofactor are likely to be fixed after they develop or to co-evolve with the ISG.

KHNYN appears to have undergone extensive evolution after ZAP emerged in tetrapods. While there could be several different evolutionary forces that led to this, changes that allow it to act as a ZAP cofactor could be responsible for some of the differences in specific residues between the bony fish and tetrapod lineages. In this study, we focused on how the CUBAN domain evolved and identified a CRM1- dependent NES present in mammalian and reptile KHNYN orthologs that is required for its nuclear-cytoplasmic shuttling, interaction with ZAP and antiviral activity. The decrease in the interaction between KHNYN and ZAP when the NES is mutated or deleted is likely at least partially due to it being sequestered in the nucleus, which leads to low levels of cytoplasmic KHNYN. However, it is possible that the nuclear-cytoplasmic trafficking pathway also makes KHNYN more competent to act as a ZAP cofactor, such as by altering its post-translational modifications or its interaction partners. The NES in KHNYN has evolved specifically in tetrapod KHNYN orthologs, with the most substantial changes in positions 3 and 4. When these positions were mutated to the residues found in zebrafish KHNYN, the protein was localized in the nucleus, did not efficiently interact with ZAP-L and had little antiviral activity. Once the full NES evolved, it became fixed in the reptile and mammal lineages.

The low abundance of KHNYN appears to be at least partly due to its CUBAN domain, which mediates an interaction with neddylated cullin-RING E3 ubiquitin ligases (36). Mutation of residues in this domain that mediate binding to NEDD8 lead to a substantial increase in KHNYN abundance, though this does not affect its antiviral activity. KHNYN protein levels could be tightly regulated to prevent off-target endoribonuclease activity, which would be determinantal to cellular gene expression, thus necessitating turnover by cullin-RING E3 ubiquitin ligases. In addition, KHNYN could have additional cellular functions beyond acting as a ZAP cofactor since KHNYN orthologs are present in bony fish, which do not have ZAP orthologs. It should be noted that the function of N4BP1 and KHNYN in fish is not known.

ZAP subcellular localization appears to regulate its antiviral activity against several viruses in that ZAP-L with an intact S-farnesylation motif mediates more potent restriction than ZAP-S (50–53). This correlates with preferential KHNYN and TRIM25 binding to ZAP-L compared to ZAP-S, even though the binding sites for KHNYN and TRIM25 are present in both isoforms (8, 20, 22, 53). KHNYN subcellular localization also appears to be important in that its CRM1 NES is required for antiviral activity. How KHNYN is targeted to the nucleus is not clear and we have not identified a canonical nuclear localization signal (NLS) in it. However, KHNYN could be trafficked into the nucleus by interacting with other proteins that contain an NLS. The relative timing for how KHNYN interacts with ZAP relative to ZAP binding to target RNA is not known. One possibility is that cytoplasmic ZAP-L molecules bind KHNYN prior to binding RNA, leading to a pre-formed antiviral complex. However, KHNYN appears to be limiting for ZAP antiviral activity because it is expressed at low levels and its overexpression potently promotes restriction of CpG-enriched HIV-1 (22, 41, 43, 45). Thus, only a small pool of ZAP molecules may be bound to KHNYN under steady-state conditions. Another possibility is that KHNYN cycles through the nucleus and cytoplasm and only interacts with ZAP after it binds RNA. This could act as a regulatory mechanism to allow endonucleolytic cleavage only for RNAs that have ZAP bound to them with a particular stoichiometry or structure. Therefore, in addition to its low abundance, nuclear localization of KHNYN could regulate its activity by preventing it from interacting with ZAP-bound RNAs that are not bona fide targets.

## MATERIALS AND METHODS

### Plasmids and Cell lines

HeLa, HEK293T and TZM-bl cells were maintained in high glucose DMEM supplemented with GlutaMAX (Thermo Fisher Scientific), 10% fetal bovine serum, 100 U/mL penicillin and 100 mg/mL streptomycin and incubated with 5% CO_2_ at 37°C. Control CRISPR and KHNYN CRISPR HeLa cells were previously described (22). HIV-1_NL4-3_ (pHIV-1_WT_) and HIV*env*_86-561_CpG (pHIV-1_CpG_) in pGL4 were previously described (22, 57). The CRISPR-resistant pKHNYN-FLAG plasmid has been previously described (22) and specific mutations were cloned into it using site-specific mutagenesis. CRISPR-resistant KHNYN-GFP constructs were made using a flexible “GGGGSGGGGSGGGG” linker between KHNYN and GFP in the context of the murine leukemia virus (MLV)-based retroviral vector MIGR1 with the GFP replaced by the Blasticidin S-resistance gene (58). Mutant KHNYN-GFP constructs were made by either synthesizing the sequence or introducing point mutations using PCR. All primers and synthesized DNA sequences were purchased from Eurofins Genomics and all PCR reactions were performed using Q5 High-Fidelity (New England Biolabs). Stable CRISPR KHNYN HeLa cells expressing CRISPR-resistant KHNYN-GFP, KHNYNmutNEDD8-GFP, KHNYNΔCUBAN-GFP, KHNYNΔNES-GFP, KHNYN-NESmut-GFP and KHNYN L676Q/L678E were produced by retroviral vector transduction.

### Transfections and infections

HeLa cells were seeded in 24-well plates at 70% confluency. Cells were transfected according to the manufacturer’s instructions using TransIT-LT1 (Mirus) at the ratio of 3 µL TransIT- LT1 to 1 µg DNA. For the HIV experiments, 0.5 µg pHIV_WT_ or pHIV_CpG_ and the designated amount of KHNYN-FLAG or GFP-FLAG for a total of 1 µg DNA were transfected. 24 hours post-transfection, the culture media was replaced with fresh media. For HIV-1_WT_ or HIV-1_CpG_ infection of HeLa cells, viral stocks were produced by co-transfecting pHIV-1_WT_ or pHIV-1_CpG_ with pVSV-G (59) into HEK293T ZAP CRISPR cells (41) and titred on TZM-bl cells.

### Analysis of protein expression by immunoblotting

48 hours post-transfection, the HeLa cells were lysed in Laemmli buffer and heated at 95°C for 10 minutes. The culture supernatant was filtered through a 0.45 µm filter and virions were pelleted by centrifugation for 2 hours at 20,000 x g through a 20% sucrose cushion in phosphate-buffered saline (PBS). Viral pellets were resuspended in 2X Laemmli buffer. Cell lysates and virion lysates were resolved on 8 to 16% Mini-Protean TGX precast gels (Bio-Rad), transferred onto nitrocellulose membranes (GE Healthcare) and blocked in 5% non-fat milk in PBS with 0.1% Tween 20. Primary antibodies were incubated overnight at 4°C followed by 3 washes in PBS with 0.1% Tween 20 and the corresponding secondary antibody was incubated for one hour. Proteins were visualized by LI-COR (Odyssey Fc) measuring secondary antibody fluorescence or using Amersham ECL Prime Western Blotting Detection reagent (GE Lifesciences) for HRP-linked antibodies with an ImageQuant (LAS8000 Mini). Primary and secondary antibodies used in this study: 1:50 HIV anti-p24Gag (60) (Mouse), 1:3000 anti-HIV gp160/120 (Rabbit, ADP421; Centralized Facility for AIDS Reagents (CFAR), 1:5000 anti-HSP90 (Rabbit, GeneTex, GTX109753), 1:1000 anti-FLAG (DYKDDDDK, Rabbit, Cell Signaling, 14793), 1:2000 anti-β-actin (Mouse, Abcam; Ab6276), 1:5000 anti-ZAP (Rabbit, Abcam, ab154680), 1:1000 anti-GFP (Mouse. Roche 11814460001), 1:5000 anti-rabbit HRP (Cell Signaling Technology, 7074), 1:5000 anti-mouse HRP (Cell Signaling Technology, 7076), 1:5000 anti-mouse IRDye 680RD (LI-COR, 926–68070), 1:5000 anti-rabbit IRDye 800CW (LI-COR, 926–32211).

### TZM-bl infectivity assay

The TZM-bl indicator cell line was used to quantify the amount of infectious virus (61–63). Briefly, cells were seeded in 24-well plates and infected by incubation with virus stocks. 48 hours post-infection, the cells were lysed and infectivity was measured by β-galactosidase expression using the Galacto-Star System following manufacturer’s instructions (Applied Biosystems). β-galactosidase activity was quantified as relative light units per second using a PerkinElmer Luminometer.

### Immunoprecipitation assays

HeLa cells stably expressing wild-type KHNYN-GFP wild-type or mutant versions were seeded in 6-well plates for 24 hours prior to immunoprecipitation. The cells were lysed on ice in lysis buffer (0.5% NP-40, 150 mM KCl, 10 mM HEPES pH 7.5, 3 mM MgCl2) supplemented with complete Protease inhibitor cocktail tablets (Sigma-Aldrich). The lysates were incubated on ice for 1 hour and centrifugated at 20,000 x g for 15 minutes at 4°C. 50 µl of the post-nuclear supernatant was saved as the input lysate and 450 µl was incubated with 5 µg of anti-GFP antibody (Roche 11814460001) for 1 hour at 4°C. Protein G Dynabeads (Invitrogen) were then added and incubated overnight at 4°C with rotation. The lysates were washed four times with wash buffer (0.05% NP-40, 150 mM KCl, 10 mM HEPES pH 7.5, 3 mM MgCl_2_) before the bound proteins were eluted with 2X Laemmli buffer and boiled for 10 minutes. Protein expression was analyzed by western blotting as described above.

### Microscopy

HeLa cells stably expressing wild-type KHNYN-GFP or versions with specific mutations were seeded in pre-treated 24-well plates 24 hours prior to immunostaining. The cells were fixed with 4% paraformaldehyde for 20 minutes at room temperature, washed once with 1X PBS, washed once in 10 mM glycine and then permeabilized for 15 minutes in 1% BSA and 0.1% Triton-X in PBS. Rabbit anti-ZAP (1:500) or rabbit anti-TRIM25 (1:500) antibodies were diluted in 1X PBS/0.01% Triton-X and the cells were stained for 1 hour at room temperature. The cells were then washed three times in PBS/0.01% Triton-X and incubated with Alexa Fluor 594 anti-rabbit (Molecular Probes, 1:500 in 1X PBS/0.01% Triton-X) for 45 minutes in the dark. Finally, the coverslips were washed three times with 1X PBS/0.01% Triton-X100 and then mounted on slides using Prolong Diamond Antifade Mountant with DAPI (Invitrogen). For the Leptomycin B treatment experiments, HeLa cells stably expressing KHNYN-GFP wild-type or mutants were seeded in pre-treated 24-well plates 24 hours prior to a four-hour treatment with 50 nM of Leptomycin B or DMSO at 37°C. After treatment, the cells were fixed and immunostained as described above. Imaging was performed on a Nikon Eclipse Ti Inverted Microscope, equipped with a Yokogawa CSU/X1-spinning disk unit, under 100x objective and laser wavelengths of 405 nm and 564 nm. Image processing and co-localization analysis was performed with Image J (Fiji) software.

### Phylogenetic analysis of KHNYN and N4BP1 and NES prediction

Amino acid sequences for KHNYN and N4BP1 were obtained from NCBI Gene, checked manually to ensure they were full-length sequences and aligned using ClustalOmega (64). The resulting alignment file was used to infer a maximum likelihood tree in the DIVEIN web server (65) using the LG substitution model with the N4BP1-like sequences from the Californian sea hare (*Aplysia californica*), and the Crown-of-thorns Starfish (*Acanthaster planci*) as outgroups. The resulting tree was visually presented and annotated using the interactive Tree of life (iTol) (66).

KHNYN protein sequences for the ConSurf analysis (37, 38, 67) were obtained from NCBI Gene, checked manually to ensure they were full-length sequences and aligned with MUSCLE (68) to produce multiple sequence alignments (MSAs). The placental mammal MSAs contained 109 sequences, the bony fish MSA contained 72 sequences, the mammal/reptile MSA contained 131 sequences and the mammal/reptile/bony fish MSA contained 203 sequences. These bespoke MSAs were used with the ConSurf Server at https://consurf.tau.ac.il/. Either human or zebrafish KHNYN was used as the Query Sequence. The settings used were the Bayesian calculation method and ‘Best model’ evolutionary substitution model. ConSurf provided several different outputs that were then used for the analysis and specific details for these outputs can be found at https://consurf.tau.ac.il/overview.php. First, the conservation score represents the evolutionary conservation at each residue, with the lowest score indicating the most conserved position in the sequence in the context of the MSA. The scores are normalized with the average score being zero and a standard deviation of one. Second, the conservation scores above and below average are each divided into 4.5 equal intervals to produce the nine categories with the most conserved residues in category 9 and the least conserved in category 1. Third, the dominant amino acid at each position in the MSA is identified as well as the percentage of sequences with this amino acid. We determined the ratio of the percentage of the dominant amino acid at each position by dividing the percentage in the mammal/reptile MSA by the percentage in the mammal/reptile/bony fish MSA. A residue that is 100% conserved in each MSA will have a ratio of 1. Only residues in the CUBAN domain that were highly conserved in the mammal/reptile MSA (ConSurf categories 7, 8 and 9) were analyzed further.

The NES was identified using the Wregex tool with the NES/CRM1 motif and the relaxed configuration (49).

### Statistical analysis

Statistical significance was determined using unpaired two-tailed *t*-tests in GraphPad. Data are represented as mean ± standard deviation and significance was ascribed to p values < 0.05.

## ACKNOWLEDGEMENTS

We thank members of the Neil and Swanson laboratories as well as Michael Malim for helpful discussions. The following reagents were obtained through the NIH AIDS Research and Reference Reagent Program, Division of AIDS, NIAID, NIH: TZM-bl from Dr. John C. Kappes, Dr. Xiaoyun Wu and Tranzyme Inc; HIV-1 p24 Hybridoma (183-H12-5C) from Dr. Bruce Chesebro. The Antiserum to HIV-1 gp120 #20 (ARP421) was obtained from the NIBSC Centre for AIDS Reagents. These studies were funded by a Medical Research Council grant MR/S000844/1 to SJDN and CMS, a Deutsche Forschungsgemeinschaft (German Research Foundation) fellowship to DK (Project number: KM 5/1-1), a Wellcome Trust Senior Research Fellowship (WT098049AIA) to SJDN, a Royal Society/Wellcome Trust Sir Henry Dale Fellowship (206200/Z/17/Z) to CO and the Francis Crick Institute, which receives its core funding from Cancer Research UK (FC001178), the UK Medical Research Council (FC001178) and the Wellcome Trust (FC001178). MF was supported by the UK Medical Research Council (MR/R50225X/1) and was a King’s College London member of the MRC Doctoral Training Partnership in Biomedical Sciences. This work was supported by the Department of Health via a National Institute for Health Research Comprehensive Biomedical Research Centre award to Guy’s and St. Thomas’ NHS Foundation Trust in partnership with King’s College London and King’s College Hospital NHS Foundation Trust.

## SUPPLEMENTAL FIGURE LEGENDS

**Fig. S1. The nuclear export signal in KHNYN is conserved in mammals, reptiles and amphibians.** Maximum likelihood phylogenetic tree of KHNYN and N4BP1 amino acid sequences. Representative sequences from mammals (light blue), reptiles (yellow), birds (light purple), amphibians (green), and bony fish (orange) were aligned and a maximum likelihood phylogeny was inferred using the LG substitution model DIVEIN. The crown-of-thorns starfish and Californian sea hare were used as outgroups to root the tree.

**Fig. S2: Regions in human, mouse and zebrafish KHNYN and N4BP1 with predicted structure by AlphaFold.** MUSCLE alignment of KHNYN and N4BP1 orthologs from human, mouse and zebrafish. The extended di-KH, UBA-like, PIN and CUBAN or CoCUN domains are marked in the alignment. Residues with a AlphaFold per-residue confidence score (pLDDT) 90>pLDDT>70 are highlighted in yellow and pLDDT>90 are highlighted in green.

**Fig. S3. There are five potential domains in bony fish KHNYN orthologs.** The AlphaFold structure confidence score (pLLDT) for zebrafish KHNYN and the ConSurf conservation score from a multiple sequence alignment of bony fish KHNYN protein sequences were plotted for each amino acid in the zebrafish KHNYN sequence.

**Fig. S4: The KHNYN CUBAN domain is essential for KHNYN antiviral activity on viral gene expression and may control its low expression level. (A)** Representative western blots for Gag and Env in cell lysates as well as virion production for the experiments shown in Figure 3B. **(B)** Expression of ZAP cofactors that regulate viral RNA degradation in HeLa cells (16-22). Data is from Nagaraj et al 2011 (43).

**Fig. S5. Characterization of KHNYN-GFP cell lines. (A)** Control CRISPR, ZAP CRISPR, KHNYN CRISPR or KHNYN CRISPR + KHNYN-GFP cells were infected with VSV-G pseudotyped HIV-1_WT_ or HIV-1_CpG_. 48 hours post-infection, the cell supernatant was harvested and infectious virus production was measured in TZM-bl cells. Each bar shows the average value of three independent experiments normalized to the value obtained for wild-type HIV-1 in control CRISPR cells. *p < 0.05 as determined by a two-tailed unpaired t-test comparing HIV-1_CpG_ in each cell line to the control CRISPR cell line. **(B)** Representative western blots for Gag and Env in cell lysates as well as virion production for the experiments shown in Figure 3C.

**Fig. S6. The nuclear export signal at the C-terminus of the KHNYN CUBAN domain is required for antiviral activity on viral gene expression. (A)** Representative western blots for Gag and Env in cell lysates as well as virion production for the experiments shown in Figure 5C. **(B)** Confocal microscopy images for wild-type KHNYN-GFP, KHNYNΔCUBAN-GFP, KHNYN-ΔNES and KHNYN- NESmut-GFP (green) co-staining with endogenous TRIM25 (magenta), scale bar is 10 µm.

**Fig. S7. The CUBAN domain is conserved in cartilaginous fish, lamprey, lancelet, echinoderm and mollusc N4BP1-like proteins.** MUSCLE alignment of CUBAN domains from KHNYN, N4BP1 or N4BP1-like proteins from the indicated species. Residues highlighted are highly conserved in the KHNYN CUBAN domain (ratio >0.85) in Figure 5A. *Amblyraja radiata*, *Callorhinchus milii*, *Carcharodon carcharias*, *Chiloscyllium plagiosum* and *Scyliorhinus canicular* are cartilaginous fishes. *Petromyzon marinus* Is a lamprey. *Branchiostoma belcheri* Is a lancelet. *Acanthaster planci* is an echinoderm. *Aplysia californica*, *Biomphalaria glabrata* and *Pomacea canaliculate* are molluscs.

**Fig. S8. L676 and F678 in the KHNYN CUBAN domain NES are required for antiviral activity on viral gene expression. (A)** Representative western blots for Gag and Env in cell lysates as well as virion production for the experiments shown in Figure 6C.

